# Volatile hydrogen cyanide released by *Pseudomonas aeruginosa* provides a competitive advantage over *Staphylococcus aureus* in biofilm and *in vivo* lung environments

**DOI:** 10.1101/2022.03.24.484598

**Authors:** Sylvie Létoffé, Yongzheng Wu, Sophie E Darch, Christophe Beloin, Marvin Whiteley, Lhousseine Touqui, Jean-Marc Ghigo

**Affiliations:** Institut Pasteur, Université de Paris, CNRS UMR 6047, Genetics of Biofilms Laboratory, 75015 Paris, France; Institut Pasteur, Université de Paris, CNRS UMR3691, Cellular Biology of Microbial Infection Laboratory, 75015 Paris, France; Department of Molecular Medicine, University of South Florida, 12901 Bruce B. Downs Boulevard, Tampa, Florida, 33612, United States; School of Biological Sciences, Georgia Institute of Technology, Atlanta, Georgia, USA; Institut Pasteur, Université de Paris, Mucoviscidose et Bronchopathies Chroniques, 75015 Paris, France; Centre de Recherche Saint-Antoine, CRSA, Sorbonne Université, Inserm, 75012 Paris, France

**Keywords:** Volatile compounds, *Staphylococcus aureus*, *Pseudomonas aeruginosa*, bacterial co-infection, hydrogen cyanide

## Abstract

Diverse bacterial volatile compounds alter bacterial stress responses and physiology, but their contribution to population dynamics in polymicrobial communities is not well known. In this study, we showed that airborne volatile hydrogen cyanide (HCN) produced by a wide range of *Pseudomonas aeruginosa* clinical strains leads to at-a-distance inhibition of the growth of a wide array of *Staphylococcus aureus* strains. We determined that low oxygen level environments not only enhance *P. aeruginosa* HCN production but also increase *S. aureus* sensitivity to HCN, which impacts *P. aeruginosa-S. aureus* competition in microaerobic *in vitro* mixed biofilms as well as in an *in vitro* cystic fibrosis lung sputum medium. Consistently, we demonstrated that production of HCN provides a competitive advantage to *P. aeruginosa* in a mouse model of airways co-infected by *P. aeruginosa* and *S. aureus*. Our study therefore demonstrates that *P. aeruginosa* HCN contributes to local and distant airborne competition against *S. aureus* and potentially other HCN-sensitive bacteria in contexts relevant to cystic fibrosis and other polymicrobial infectious diseases.

**IMPORTANCE:** Airborne volatile compounds produced by bacteria are often only considered as attractive or repulsive scents, but they also directly contribute to bacterial physiology. Here we showed that volatile hydrogen cyanide (HCN) released by a wide range of *Pseudomonas aeruginosa* clinical strains inhibits *Staphylococcus aureus* growth in low oxygen *in vitro* biofilms or aggregates and *in vivo* lung environments. These results are of pathophysiological relevance, since lungs of cystic fibrosis patients are known to present microaerophilic areas and to be commonly associated with the presence of *S. aureus* and *P. aeruginosa* in polymicrobial communities. Our study therefore provides insights into how a bacterial volatile compound can contribute to the exclusion of *S. aureus* and other HCN-sensitive competitors from *P. aeruginosa* ecological niches. It opens new perspectives for the management or monitoring of *P. aeruginosa* infections in lower lung airway infections and other polymicrobial disease contexts.

## INTRODUCTION

Bacteria release a wide diversity of volatile molecules contributing to cross-kingdom interactions with fungi, plants and animal (1, 2). Bacteria volatile compounds also play a role in bacterial physiology by altering stress responses, antibiotic resistance, biofilm formation and expression of virulence factors. Although these interactions likely contribute to bacterial population dynamics, relatively little is known regarding interactions mediated by volatile compounds in polymicrobial communities (2–8). Cystic Fibrosis (CF) is a common genetic disease in which the patients’ airways are often colonized by multiple bacterial pathogens, including *Pseudomonas aeruginosa* and *Staphylococcus aureus*, that are frequently found in association in the same lung lobes (9–15). Whereas *S. aureus* usually colonizes the airways first during CF infection, it is later outcompeted and replaced by *P. aeruginosa* (12, 16–20).

Several *P. aeruginosa* extra-cellular factors inhibiting *S. aureus* growth could contribute to this colonization shift during CF infection, including siderophores, 4-hydroxy-2-heptylquinoline-*N*-oxide (HQNO), proteases, RedOx and surface active compounds (19, 21–26). By contrast, less is known about how competitive interactions of *P. aeruginosa* are mediated via the production of volatile compounds and their impact on the dynamics of coinfections with *S. aureus* (27–29).

It has long been recognized that *P. aeruginosa* metabolism produces volatile hydrogen cyanide (HCN) that can rapidly diffuse into the environment (30, 31). HCN is an inhibitor of cytochrome c oxidases and other metallo-enzymes that bind iron, leading to the inhibition of the respiratory chain (30). HCN production is restricted to *Pseudomonas, Chromobacterium, Rhizobium* and several cyanobacterial species that avoid auto-intoxication by expressing HCN-insensitive cytochrome oxidase (31). *P. aeruginosa* HCN is produced by the oxidative decarboxylation of glycine mediated by membrane-bound cyanide synthases encoded by the *hcnABC* operon (32–34). *hcnABC* expression is maximal between 34 °C and 37 °C and transcriptionally up-regulated in microaerophilic conditions or by high bacterial cell density conditions (31, 35). Consistently, HCN production by *P. aeruginosa* is regulated by the anaerobic regulator Anr, and the LasR and RhlR quorum sensing regulators (36). HCN is therefore produced in environmental conditions leading to the induction of *P. aeruginosa* virulence factors, including the synthesis of alginate a constituent of *P. aeruginosa* biofilms matrix and a major virulence factor in the lungs of CF patients (37–39).

Considering that HCN was shown to poison a wide range of eukaryotic organisms (2, 40–42), it was hypothesized that cyanogenesis could also poison HCN-sensitive bacteria in a range of polymicrobial niches (26, 28, 30, 31, 39, 43, 44). However, whereas *P. aeruginosa* HCN was shown to inhibit the growth of a wide range of Staphylococci, including *S. aureus* (45), the direct contribution of HCN to *P. aeruginosa* dominance over *S. aureus* within polymicrobial niches such as biofilms or infected lungs is still unclear.

Here we showed that exposure to airborne HCN produced by *P. aeruginosa* inhibits *S. aureus* growth and influences the dynamics of *P. aeruginosa*-*S. aureus* interactions in *in vitro* mixed biofilms. We determined that HCN production is widespread among *P. aeruginosa* clinical strains and particularly active in low oxygen (microaerobic) conditions against a representative panel of *S. aureus* isolates. We also demonstrated that *P. aeruginosa* HCN impairs *S. aureus* growth in an *in vitro* CF lung sputum model as well as in a mouse model of airway co-infection by *P. aeruginosa* and *S. aureus*. Our study therefore shows that volatile HCN provides *P. aeruginosa* with a competitive advantage in local and at-a-distance airborne competitions against *S. aureus* and potentially other HCN-sensitive bacteria in context relevant to CF and other polymicrobial infectious diseases.

## RESULTS

### Production of volatile hydrogen cyanide by *Pseudomonas aeruginosa* leads to airborne inhibition of *Staphylococcus aureus* growth

To determine whether HCN released by *P. aeruginosa* could inhibit the growth of *S. aureus*, we first tested its HCN production by PAO1, a commonly used strain of *P. aeruginosa* isolated from a wound infection **(46)**. Using a semi-quantitative HCN detection method based on the intensity of blue color produced upon HCN reaction with copper(II) ethylacetoacetate and 4,4’-methylenebis- (N,N-dimethylaniline) (47) (Supplementary Fig. S1), we detected an HCN signal emitted from WT *P. aeruginosa* PAO1 grown in LB, which increased upon glycine supplementation (Fig.1A). By contrast, no HCN signal could be detected from a Δ*hcnB* mutant, which lacks HCN production (Fig.1A). We then exposed *S. aureus* to *P. aeruginosa* PAO1 volatile compounds in the set-up described in the supplementary Fig. S1. Whereas exposure to *P. aeruginosa* PAO1 cultures modestly reduced *S. aureus* MW2 growth, the aerial exposure to culture supplemented with glycine led to a 100-1000-fold growth inhibition dependent on *hcnB* (Fig.1B). Moreover, we observed that a *S. aureus* MW2 *srrAB* mutant lacking the SrrAB global regulator of the transition from aerobic to anaerobic respiration displayed an increased sensitivity to HCN (Supplementary Fig. S2) (48). Finally, preventing HCN release by placing a parafilm seal on the emitting plate containing *P. aeruginosa* or *P. aeruginosa ΔhcnB* culture supplemented with glycine did not lead to any growth defect, confirming the contribution of volatile HCN to *S. aureus* MW2 growth inhibition (Fig.1B).

**Figure 1.**
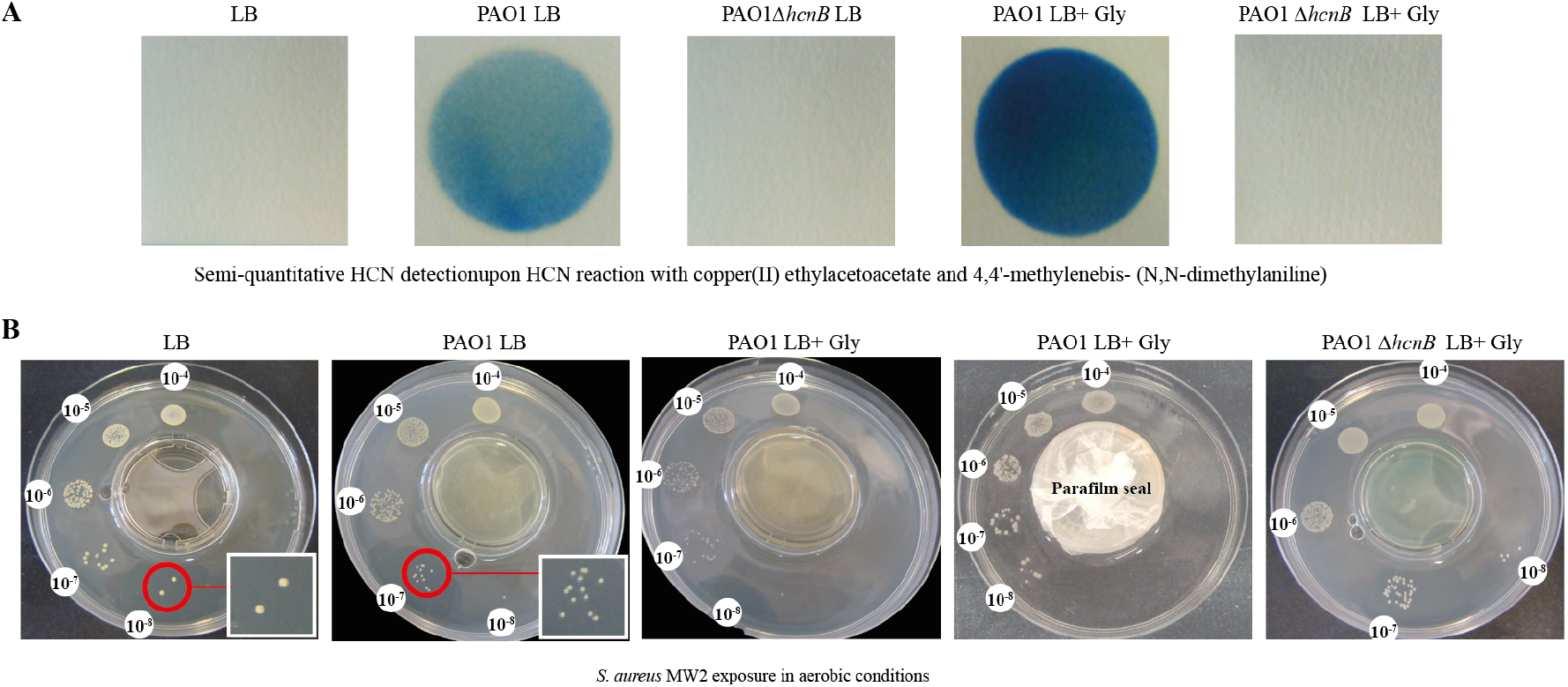
*P. aeruginosa* production of hydrogen cyanide leads to airborne inhibition of *S. aureus* growth. **A:** Semi-quantitative detection of volatile HCN emitted from *P. aeruginosa* WT and Δ*hcnB*, in LB supplemented or not with 0.4% (w/v) glycine, after 24h incubation at 37°C in aerobic conditions. HCN detection is based on HCN reaction with copper (II) ethylacetoacetate and 4,4′-methylenebis-(N,N-dimethylaniline). **B:** Serial dilution of *S. aureus* MW2 upon exposure to *P. aeruginosa* WT or Δ*hcnB* cultures in LB supplemented or not with 0.4% (w/v) glycine in the 2-Petri-dish assay (see Supplementary Fig. S1). No inhibition of *S. aureus* MW2 growth is observed when the middle plate containing *P. aeruginosa* culture is covered and sealed with parafilm. Pictures were taken after 24h of incubation at 37°C in aerobic conditions. Each experiment was performed at least three times.

### Production of biogenic HCN is widespread among *Pseudomonas aeruginosa* clinical strains and active against diverse *S. aureus* isolates

To determine whether HCN production is a widespread *P. aeruginosa* property, we exposed the HCN-sensitive *S. aureus srrAB* mutant to a panel of laboratory and clinical *P. aeruginosa* strains, many of them isolated from airway infections (Table 1). We showed that, despite variations, all tested strains aerially inhibited *S. aureus srrAB* in aerobic conditions, even in the absence of glycine (Supplementary Fig. S3A). Moreover, *P. aeruginosa* strains that led to a minimal reduction of *S. aureus* growth showed a strong growth inhibition phenotype when grown in the presence of glycine, indicative of an increase in HCN production (Supplementary Fig. S3B). In addition, we also showed that the production of HCN by *P. aeruginosa* PAO1 in the presence of glycine inhibited the growth of a wide panel of distinct pathogenic *S. aureus* strains (Supplementary Fig. S4), confirming HCN-mediated growth inhibition at distance over a broad range of *S. aureus* strains.

**Table 1:** Plasmids and strains used in the study.

|  | Relevant characteristics or origin | Reference / source |
| --- | --- | --- |
| **Plasmids** |  |  |
| pMRP9-1 | *gfp*-expressing plasmid | (70) |
| ***Pseudomonas aeruginosa* strains** |  |  |
| PAO1 | Isolate from wound infection, Melbourne | (46) |
| PAO1*gfp* | Green fluorescent PAO1 containing pMRP9-1 | This study |
| PAO1 $\Delta hcnB$ | HCN deficient $\Delta hcnB$ strain | University of Washington<br>Genome Center, Gift from<br>O. Lesouhaitier |
| PAO1 $\Delta hcnB$ *gfp* | Green fluorescent PAO1$\Delta hcnB$ containing pMRP9-1 | This study |
| PA14 | Clinical isolates from burn patients | (71) |
| PAK | Virulent strain sensitive to Pf phage | (72) |
| 7508 | Bronchial secretion | (73) |
| 8931 | Lung transplant | (73) |
| 9854 | Nasal swab | (73) |
| 11989 | Tonsil swab | (73) |
| 12269 | Sputum | (73) |
| 13305 | Bronchial secretion | (73) |
| Psae1152 | Drainage catheter | (73) |
| Psae1471 | Respiratory tract | (73) |
| Psae1659 | Respiratory tract | (73) |
| Psae1716 | Blood | (73) |
| Psae1928 | Respiratory tract | (73) |
| Psae2328 | Urine | (73) |
| BJN8 | Catheter related blood stream infection related BSI from Beaujon Hospital. Patient#8. | (74) |
| BJN33 | Catheter related blood stream infection related BSI from Beaujon Hospital. Patient#33. | (74) |
| BJN53 | Catheter related blood stream infection related BSI from Beaujon Hospital. Patient#53. | (74) |
| BJN66 | Catheter related blood stream infection related BSI from Beaujon Hospital. Patient#66. | (74) |
| ***Staphylococcus aureus*** |  |  |
| MW2 | Community-acquired methicillin-resistant | (75) |
| MW2$\Delta srrAB$ | Deletion of *srrAB* genes | Gift from I. Lasa (76) |
| 15981 | Biofilm-forming strain isolated at the Microbiology Department of the University Clinics of Navarra | (77) |
| COL | Initially isolated from the operating theatre in a hospital in Colindale, England in the early 1960s. | (78) |
| Newman | *S. aureus* strain Newman was isolated in 1952 from a human infection | (79) |
| Xen36 | Xen 36 | Caliper Life Sciences, |
| HG001 | Highly virulent strain derivative of NCTC 8325 originally used to propagate bacteriophage 47 | (80) |
| N315 | hospital-acquired methicillin-resistant isolated in 1982 from a pharyngeal smear of a patient in Japan. | (81) |
| USA300LAC | Epidemic community-associated methicillin-resistant isolated from the Los Angeles County jail | (82) |
| USA300LAC*dsred* |  | (83) |
| V329 | Biofilm forming bovine mastitis isolate | (84) |

### Microaerobia enhances *P. aeruginosa* HCN production while increasing *S. aureus* sensitivity to HCN

*P. aeruginosa* HCN production is regulated by quorum sensing (36). Consistently, we observed an increase of the HCN signal during the transition from exponential to stationary phase, at culture densities OD_600_>2 (Fig. 2A). Compared to aerobic conditions, we also observed that the *P. aeruginosa* HCN signal was enhanced in microaerobic conditions (0.4-0.8% O_2_) (Fig. 2BD) (33). Moreover, in these microaerobic conditions, *S. aureus* MW2 was more sensitive to *P. aeruginosa* PAO1 HCN that when grown in aerobic conditions (Fig. 2CE). These results suggest that microaerobic conditions not only lead to higher HCN production in *P. aeruginosa* PAO, but also increased *S. aureus* MW2 sensitivity to HCN.

**Figure 2.**
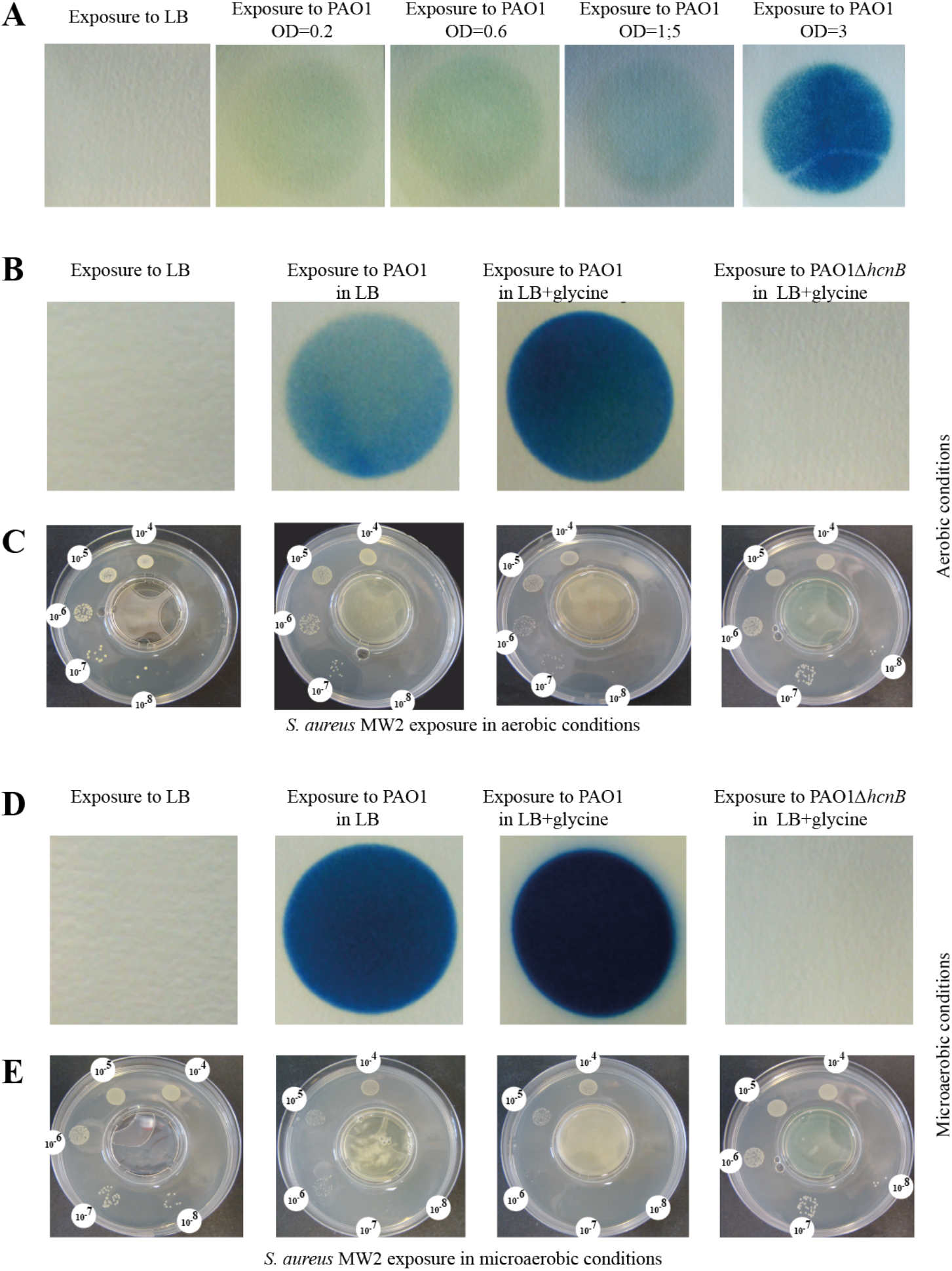
Microaerobia stimulates *P. aeruginosa* HCN production and increase *S. aureus* sensitivity to HCN. **A:** Semi-quantitative detection of volatile HCN showed an increased production of HCN by *P. aeruginosa* PAO1 at different stages of growth stages in LB medium. Each experiment was performed at least three times. **B, D:** Semi-quantitative detection of volatile HCN from *P. aeruginosa* WT and Δ*hcnB*, in LB supplemented or not with 0.4% (w/v) glycine, after 24h incubation at 37°C in aerobic (B) or microaerobic (D) conditions. Each experiment was performed at least three times. **C,E:** Growth of serial dilution of *S. aureus* MW2 WT upon exposure to *P. aeruginosa* WT or Δ*hcnB* cultures in LB supplemented or not with 0.4% (w/v) glycine, after 24h incubation at 37°C in aerobic (C) or microaerobic (E) conditions, using the 2-petri-dish assay described in Fig S1. Each experiment was performed at least three times.

### Production of biogenic HCN impairs *S. aureus* growth in *in vitro* mixed biofilms

Our results suggested that HCN production could contribute to the dynamics of *P. aeruginosa-S. aureus* competition. Considering that microaerobic conditions prevail within multi-species biofilms (49), we hypothesized that *P. aeruginosa* HCN production in biofilms could impact *S. aureus* growth dynamics in mixed *P. aeruginosa / S. aureus* biofilms. To test this *in vitro*, we co-inoculated continuous-flow biofilm microfermenters with *P. aeruginosa* PAO1 WT (HCN+) and or Δ*hcnB* (HCN-) mutant at a 1:1 ratio with three different *S. aureus* strains, including HG001, or Xen36 and MW2. While all strains displayed similar individual biofilm-forming capacities (Fig. 3A), the *P. aeruginosa* and *S. aureus* proportions in the resulting two-species biofilms formed after 48h showed that all tested *S. aureus* strains formed less biofilm biomass, as measured by CFU count, when mixed with WT *P. aeruginosa* than with the HCN-deficient mutant (Fig. 3B). Taken together, these results indicate that the production of *P. aeruginosa* biogenic HCN impairs growth and outcompete *S. aureus* in mixed biofilms.

**Figure 3.**
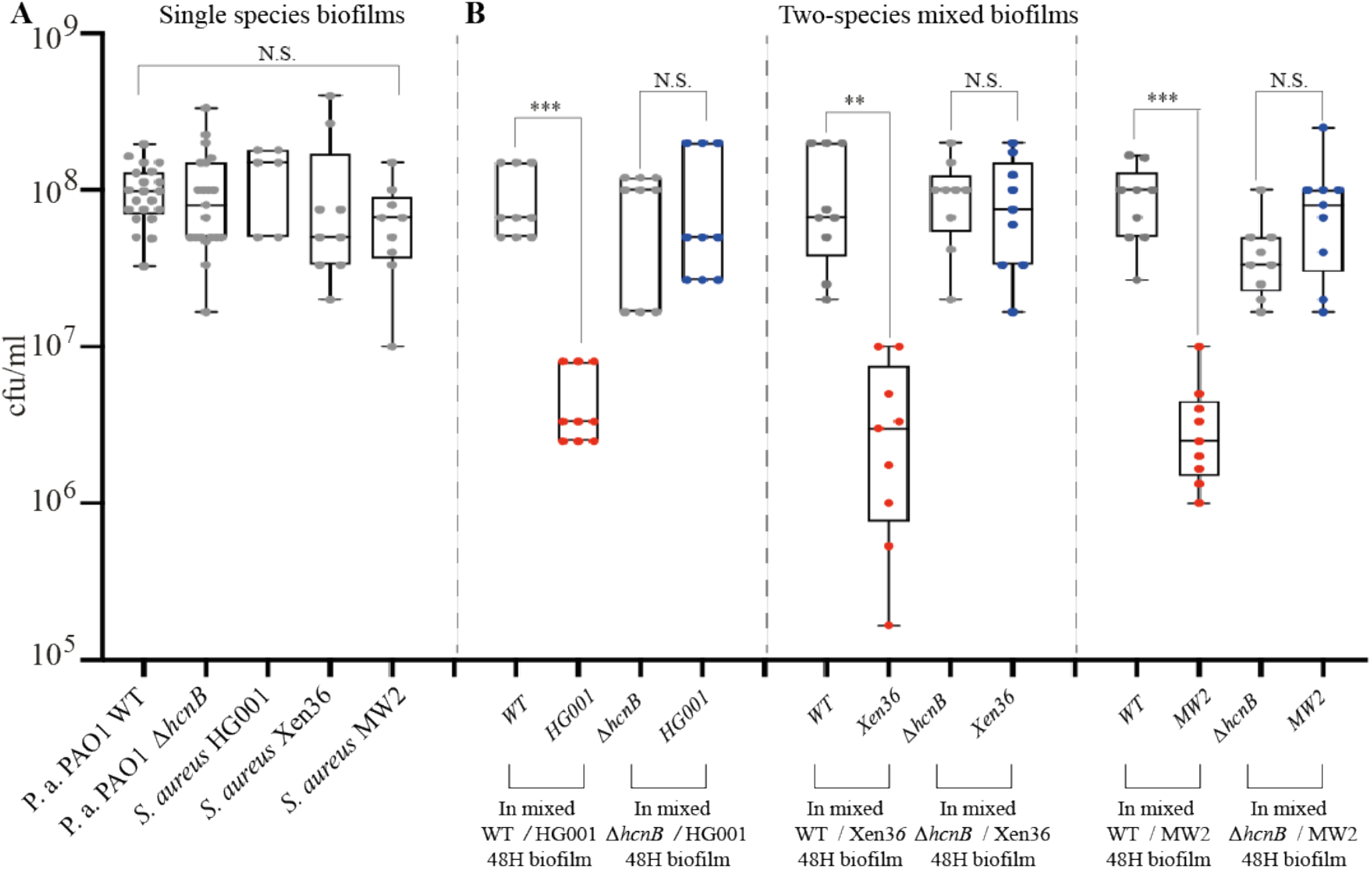
Production of biogenic HCN impairs *S. aureus* growth in *in vitro* mixed biofilms. **A:** Number of CFU in indicated single species biofilms grown in continuous-flow microfermenters, in LB medium for 48h at 37°C. **B:** Number of CFU in indicated two-species mixed biofilms: each *S. aureus* strain was mixed with either WT *P. aeruginosa* PAO1 or its Δ*hcnB* mutant at a 1:1 ratio. The biofilms were grown in continuous flow microfermenters in LB medium for 48h at 37°C. Statistics correspond to two-tailed unpaired *t-test* with Welch correction. N.S.: not significant, ** p≤0.01 and *** p≤0.001.

### Production of biogenic HCN provides a competitive advantage to *P. aeruginosa* over *S. aureus* in CF-relevant conditions

To test whether production of HCN could provide *P. aeruginosa* with a competitive advantage over *S. aureus* in CF-relevant conditions, we first used the synthetic CF sputum (SCFM2) medium, designed to recapitulate human CF environments (50). We inoculated this medium with red fluorescent *S. aureus LACdsrfp* alone or in 1:1 mix ratio with *P. aeruginosa PAO1gfp* WT (HCN+) or its Δ*hcnB* (HCN-) mutant. The comparison of the respective *S. aureus* and *P. aeruginosa* spatial organization revealed a strong reduction of *S. aureus* biomass development (Fig. 4A) and aggregate abundance when co-inoculated with WT *P. aeruginosa* (Fig. 4B top), in contrast to the opposite increase of *S. aureus* development in presence of the *P. aeruginosa ΔhcnB* mutant (Fig. 4B bottom).

**Figure 4.**
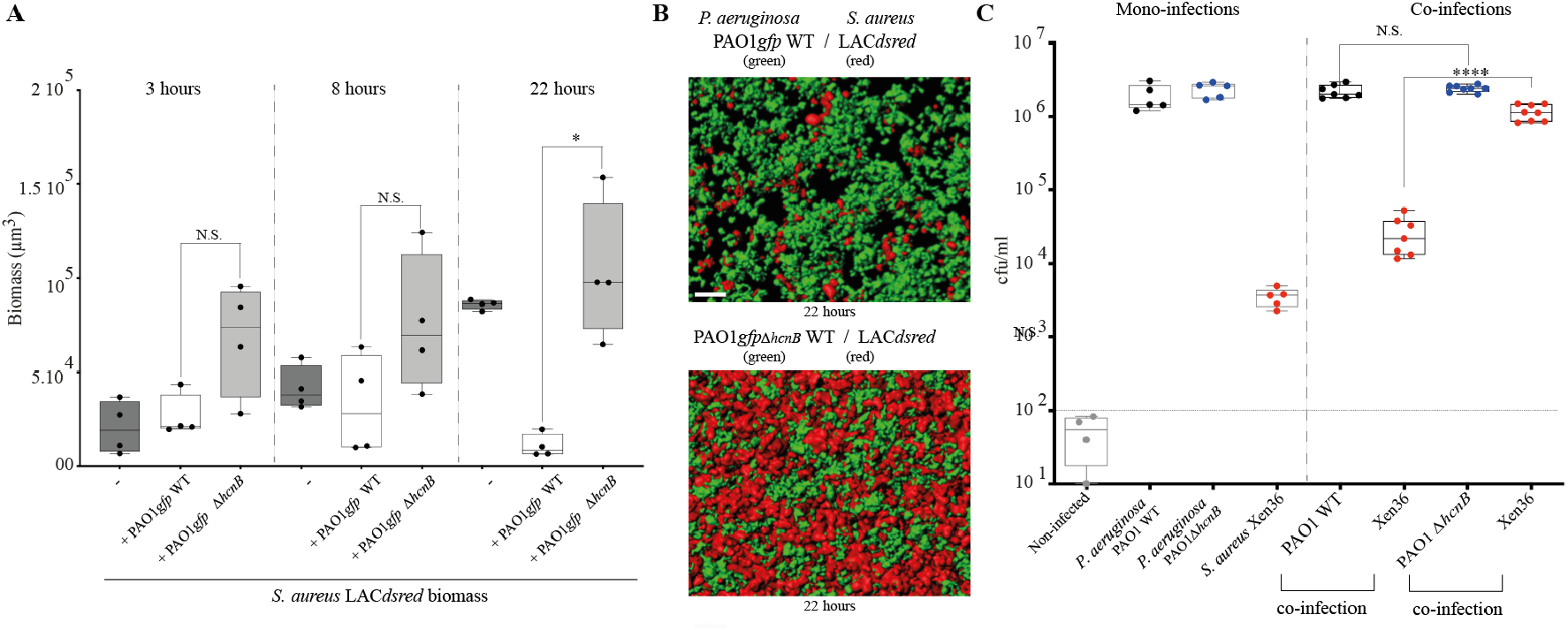
*P. aeruginosa* production of biogenic HCN inhibits *S. aureus* growth in in CF-relevant conditions. **A:** Total biomass of *S. aureus* aggregates as monoculture or in co-infection with PAO1 wildtype and/or *hcnB* mutant. Isolates were cultured in SCFM2 and imaged using confocal microscopy at 3, 8 and 22 hours. Statistics correspond to two-tailed unpaired *t-test* with Welch correction. N.S.: not significant, * p≤0.05. **B:** Representative rendered confocal micrograph of *S. aureus* and *P. aeruginosa* coinfection in SCFM2. Top – wild type *P. aeruginosa* in green, *S. aureus* in red. Bottom – *P. aeruginosa hcnB* mutant in green, *S. aureus* in red. **C**: *In vivo* competition experiments in mice lungs. Mono and 1:2 mixed ratio co-inoculation of *S. aureus* Xen36, *P. aeruginosa* PAO1 WT or Δ*hcnB* mutant. Number of bacteria CFU was counted in the lung homogenates of mice 24 h after infection. Non infected SOPF mice showed minimal lung bacterial contamination with CFY<100 (horizontal dotted line). Statistics correspond to two-tailed unpaired *t-test* with Welch correction. N.S.: not significant, *** p≤0.0001.

To further test the *in vivo* impact of HCN production on *S. aureus/P. aeruginosa* mixed community dynamics in the microaerophilic lung airways, we performed an *in vivo* competition in mice, in which lungs were intratracheally co-inoculated with *S. aureus* Xen36 strain and either *P. aeruginosa* PAO1 WT or Δ*hcnB*. Mice lungs infected individually show that *P. aeruginosa* colonizes better than *S. aureus* Xen36 (Fig. 4C left) The comparison of the number of CFUs extracted from mice lungs co-inoculated in a 2:1 mixed ratio (*P. aeruginosa: S. aureus)* showed a 2 log decrease of *S. aureus* Xen36 in presence of WT PAO1 (HCN+) as compared to *S. aureus* growth in presence of the PAO1 Δ*hcnB* mutant (HCN-) (Fig.4C). Taken together, our results demonstrate that HCN production by *P. aeruginosa* reduces *S. aureus* colonization in co-infected mouse lungs and other microaerobic biofilm-like environments.

## DISCUSSION

In this study, we showed that airborne HCN produced by a wide range of *P. aeruginosa* clinical strains is enhanced in microaerobic conditions and inhibits various *S. aureus* isolates *in vitro*, disadvantaging *S. aureus* colonization in polymicrobial biofilms. This occurs both in a CF sputum medium and in an *in vivo* mouse model of pulmonary co-infection. A number of bacterial infections are characterized by the development of polymicrobial communities in which complex interactions between bacteria can influence the outcome of diseases (12, 17, 43, 51). Colonization of the lungs during CF is one of the best examples of polymicrobial infection that is characterized by excessive mucus production in airways and decreased mucosal clearance (52). This favors lung colonization by bacterial pathogens, including *P. aeruginosa*, *S. aureus*, non-typeable *Haemophilus influenzae* and *Burkholderia cepacia*, where prevalence varies with the age and treatments of CF patients (52–55). These airway infections are difficult to eradicate despite aggressive antibiotic therapy (56, 57) and are associated with inflammation, leading to a progressive decline in lung functions and, ultimately, to respiratory failure (53, 57–59).

Interactions between microorganisms have been shown to be key determinants of their distribution and activity in most ecosystems (12, 58) with several *P. aeruginosa* secreted molecules shown to inhibit *S. aureus*’s growth (16, 19, 25, 26, 60). By contrast to local competition driven by short range diffusion (<10 μm) of most inhibitory metabolic products, volatile HCN produced by *Pseudomonas* and a number of bacterial species could play an important role in the spatial organization of microbial communities (31). HCN could indeed contribute to both local and distant, airborne competition between microorganisms in physically heterogeneous solid, liquid and gaseous environments such as the lungs and other organic tissues.

Our results are of pathophysiological relevance, since CF lungs are known to present microaerophilic areas and to be commonly associated with the presence of *S. aureus* and *P. aeruginosa* multispecies biofilms, reaching high *P. aeruginosa* cell density, two conditions that have been shown to induce *hcnABC* gene expression and subsequent HCN production (30, 31, 34, 36, 61). HCN was indeed previously detected in the sputum and bronchial-alveolar lavage fluids of CF patients infected by *P. aeruginosa* and the measure of HCN levels in lungs of CF patients has been used as a non-invasive breath test to diagnose *P. aeruginosa* infection (44, 62–64). This suggests that the levels of HCN produced by *P. aeruginosa* in the lung environment could be sufficient to poison aerobic metabolism and growth, excluding competitors from *P. aeruginosa* ecological niches (39, 44).

*S. aureus* had been regarded as one of the initial microbial colonizers of the CF patients’ airways before being displaced by *P. aeruginosa* (14, 20) and our results support the hypothesis that metabolic poisoning upon HCN production could be a key determinant of *Pseudomonas* distribution in the lung upon exclusion of *S. aureus* in mixed *in vitro and in vivo* polymicrobial biofilms (19, 28, 34, 39). However, *P. aeruginosa* PAO1 was also shown to reduce its toxicity towards *S. aureus* or to facilitate *S. aureus* microcolony formation through alginate production, therefore promoting the coexistence of these two bacteria (13, 65–67). Consistently, we observed that, although production of HCN by PAO1 WT reduced the number of *S. aureus* recovered from co-inoculated lungs compared to co-inoculation with PAO1Δ*hcnB*, there was a 1000-fold increase in *S. aureus* abundance when comparing mono-inoculation and co-inoculation with PAO1Δ*hcnB* (Fig. 4C). This indicates that, in absence of HCN, *S. aureus* growth is stimulated by *P. aeruginosa*, which further emphasizes that these two bacteria could engage in complex negative and positive interactions *in vivo* (67, 68).

Our study therefore contributes to a better understanding of *P. aeruginosa* and *S. aureus* competition in a context relevant to CF airway infection. Whereas further studies will be required to tease out the respective ecological contribution of HCN and other *P. aeruginosa* factors to outcompete *S. aureus*, our results further illustrate the remarkable ability of *P. aeruginosa* to adapt and thrive in multispecies communities. The identification of a volatile compound-based mechanism potentially underlying the dynamic shift from *S. aureus* to *P. aeruginosa* dominance in polymicrobial infection opens new perspectives for the management or monitoring of *P. aeruginosa* infections in lower lung airway infections and other polymicrobial disease contexts.

## MATERIALS AND METHODS

### Bacterial strains, plasmids and growth conditions

Bacterial strains and plasmids used in this study are listed in Table 1. All experiments were performed in lysogeny broth (LB) medium, supplemented or not with 0.4% (w/v) glycine and incubated at 37°C. All chemicals were purchased from Sigma-Aldrich.

### Test in SCFM2 artificial sputum model

Green fluorescent *P. aeruginosa* (WT or HCN mutant) carrying pMRP9-1 and *S. aureus* expressing dsRed red fluorescent protein (see Table 1) were grown overnight in Tryptic Soy Broth (TSB). Cells were washed twice and resuspended in PBS. The optical density at 600 nm (OD_600_) was measured with a spectrophotometer, and washed bacterial cultures were inoculated into SCFM2 at an OD_600_ = 0.05 (~10^7^ CFU/mL) as individuals or in combination. Cultures were vortexed for 5 to 10 s to disperse bacterial cells in SCFM2. Five hundred microliters of inoculated SCFM2 was then transferred into each well of four-well microchamber slides (Lab-Tek; Nunc) and incubated under static conditions at 37°C.

#### Imaging

All images were acquired with Zeiss LSM 700 and LSM 880 confocal laser scanning microscopes utilizing Zen image capture software. Bacterial cells were visualized via GFP with an excitation wavelength of 488 nm and an emission wavelength of 509 nm or via dsRed with an excitation wavelength of 587 nm and an emission wavelength of 610 nm or with a 63× oil immersion objective. SCFM2 images were acquired by producing 512-by 512-pixel (0.26-by 0.26-μm pixels) 8-bit z-stack images that were 100 μm from the base of the coverslip. The total volumes of 100-μm z-stack images were 1822.5 mm^3^. Control images of uninoculated SCFM2 were acquired by using identical settings to determine the background fluorescence for image analysis.

#### Image analysis

All imaging was performed with identical image capture settings. To determine the background fluorescence in SCFM2, a histogram of detected dsRed and GFP fluorescence was produced in Imaris v 8.3.1 (Bitplane) for uninoculated SCFM2, and the average of the three highest voxel values was determined as the background fluorescence. Averaging across all of the control images, this value was then subtracted from all experimental images with Imaris. For aggregate and biomass quantification in

SCFM2, isosurfaces were produced for all remaining voxels after background subtraction with the surpass module in Imaris. To detect individual aggregates, the split objects option in Imaris was enabled and aggregates were defined as objects with volumes of >5 μm^3^. The total biomass (all voxels detected), average aggregate volume and number of aggregates were calculated within the vantage module in Imaris. Detected aggregate isosurfaces were then ordered by volume. Objects that were ≥0.5 and ≤5.0 μm^3^ were categorized as dispersed biomass, and objects that were >5.0 μm^3^ were categorized as aggregated biomass. All image data were exported into Microsoft Excel 2016, and graphs were generated with GraphPad Prism 7.

### Screening for volatile-mediated HCN phenotypes

To evaluate the activity of HCN released by bacterial liquid culture on recipient test bacteria, a lidless 3.5 cm Petri dish was placed inside a 9 cm Petri dish, which external ring was filled with 20 mL of 1.5% LB agar (Supplementary Fig. S1A) (4). Tested recipient bacteria were spotted as 20 μL drops of 10^-4^ to 10^-8^ serial dilutions of an overnight culture adjusted to OD_600_ = 1 filled and *P. aeruginosa b*acterial liquid culture releasing or not volatile HCN were adjusted to OD_600_ = 3 and introduced in the middle of an uncovered Petri dish. The large Petri dish was then closed and incubated for 24 h at 37°C, in aerobic or microaerobic conditions. Exposure under microaerobic conditions (0.4 - 0.8% O_2_) was performed in a C400M Ruskinn anaerobic-microaerophilic station.

### Detection of HCN production

Semi quantitative determination of the levels of HCN production used a method adapted from previous studies (47). Briefly, using the set-up described in Supplementary Fig. S1A, Whatman chromatography paper soaked into HCN detection reagent containing copper(II) ethyl acetoacetate (100mg) and 4,4’-methylenebis-(N,N-dimethylaniline) (100mg) solubilized in 20 mL chloroform was laid on the surface of the central, uncovered 3.5cm Petri dish of containing bacterial liquid culture releasing or not volatile HCN. The large Petri dish was then closed and incubated for 24 h at 37°C in aerobic or microaerobic conditions. Exposure under microaerobic conditions (0.4 – 0.8% O_2_) was performed in a C400M Ruskinn anaerobic-microaerophilic station. The level of HCN was evaluated based on the intensity of blue color resulting from exposure to bacterial HCN.

### In vivo experiments

Specific opportunistic pathogen free (SOPF) Balb/c mice (male, 7 weeks, in particular free of detectable *S. aureus* and *P. aeruginosa* strain) were ordered in Janvier Labs (France) and housed in the Institut Pasteur animal facilities. All experiments were approved by the Ethics Committee of Institut Pasteur (reference 2014-0014). Mice were infected intratracheally as described previously (19). In brief, mice were anesthetized by intraperitoneal injection of ketamine (Imalgene 1000^®^, 90mg/kg)/xylazine (Rompun^®^, 10mg/kg) suspended in PBS. The anesthetized animals were subjected to non-invasive intratracheal catheterization through which *P. aeruginosa* (1×10^6^ CFU) and/or *S. aureus* (5×10^5^ CFU) suspended in 50 μL of PBS was/were introduced to initiate the infection. Twenty-four hours post infection, the animals were sacrificed by intraperitoneal injection of a lethal dose of pentobarbital. The lungs were harvested and homogenized as described previously (19). The lung homogenates were serially diluted, and the number of bacterial CFU in the lung was determined by plating and counting bacteria on LB agar (all bacteria) and/or on *P. aeruginosa* selective PIA plates and *S. aureus* selective MSA plates.

### Biofilm competition experiments in microfermenters

Continuous-flow biofilm microfermenters containing a removal glass spatula were used as described in (69) (see also https://research.pasteur.fr/en/tool/biofilm-microfermenters/). Medium flow was adjusted to 60 mL/h with internal bubbling agitation with filter-sterilized compressed air to minimize planktonic growth over biofilm development. Inoculation was performed by dipping the glass spatula for 10 min in overnight OD_600_=1 LB cultures of *S. aureus* and *P. aeruginosa* strains mixed at a 1:1 ratio. The spatula was then reintroduced into the microfermenter and the resulting 48 h mixed biofilms grown on the fermenter spatula were recovered and corresponding serial dilutions were plated on LB agar (all bacteria) and/or on *P. aeruginosa* selective PIA plates and *S. aureus* selective MSA plates.

### Statistical analysis

Two-tailed unpaired *t-test* with Welch correction analyses were performed using Prism 9.0 for Mac OS X (GraphPad Software). Each experiment was performed at least three times.

## ACKNOWLEDGEMENTS

We thank Rebecca Stevick and Inigo Lasa for critical reading of the manuscript. We are grateful to O. Lesouhaitier for generously providing the *P. aeruginosa ΔhcnB* mutant and to I. Lasa, L. Debarbieux et Susanne Haussler for providing some of the *S. aureus* and *P. aeruginosa* strains used in this study.

## AUTHORS CONTRIBUTION

S.L., Y.W., S.E. D., J.-M.G and C.B. performed the experiments; J.-M.G., S.L., L.T., C.B., M.W and S.E.D designed the experiments. S.L., C.B., L.T. S.E.D, M.W and J.-M.G., analyzed the data. J.-M.G. provided resources and funding. J.-M.G. wrote the manuscript with significant contribution from all co-authors.

