## Supplementary Figures S1 to S4 for "Volatile hydrogen cyanide released by *Pseudomonas aeruginosa* provides a competitive advantage over *Staphylococcus aureus* in biofilm and *in vivo* lung environments"

### SUPPLEMENTARY MATERIALS

This PDF file includes supplementary Figures S1 to S4

### SUPPLEMENTARY FIGURES

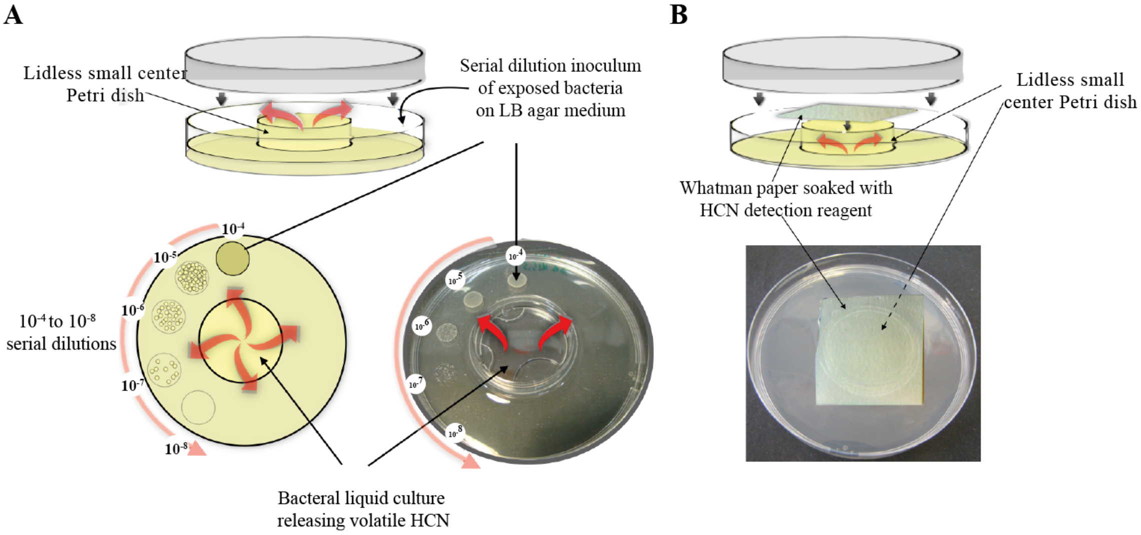


Supplementary Figure S1. **Two-Petri-dish assay. A:** Evaluation of volatile-mediated impact on growth between physically separated bacteria. A small lidless Petri dish is placed inside a larger one, which is closed by its lid. Serial dilutions of bacteria spotted on external LB agar ring are exposed to volatile molecule released from the culture placed in the central small Petri dish. Bacterial growth was monitored after 24h of incubation at 37°C, in aerobic or microaerobic conditions. **B:** Semi-quantitative HCN detection: Whatman chromatography paper soaked with HCN detection reagent was laid on the surface of the uncovered little Petri dish containing bacterial liquid culture releasing or not volatile HCN, and the large Petri dish was then closed and incubated for 24h at 37°C in aerobic or microaerobic conditions.

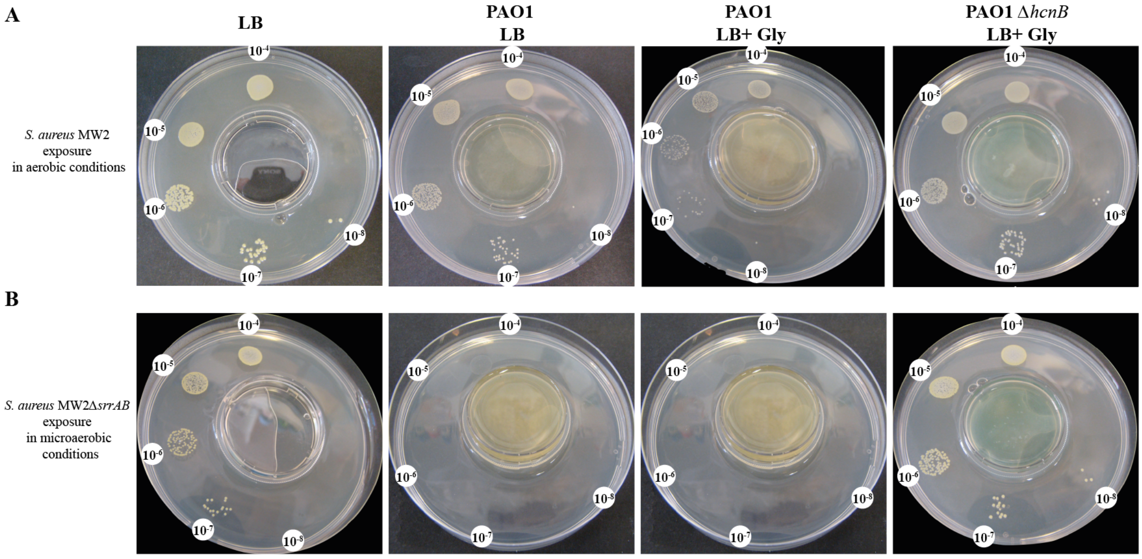


Supplementary Figure S2**. *S. aureus* MW2 *srrAB* mutant displayed increased sensitivity to HCN.** Growth of serial dilution of *S. aureus* MW2 WT (A) or *srrAB* mutant (B) upon exposure to *P. aeruginosa* WT or Δ*hcnB* cultures in LB supplemented or not with 0.4% (w/v) glycine, after 24h incubation at 37°C in aerobic conditions, using the 2-petri-dish assay described in Fig S1. Each experiment was performed at least three times.

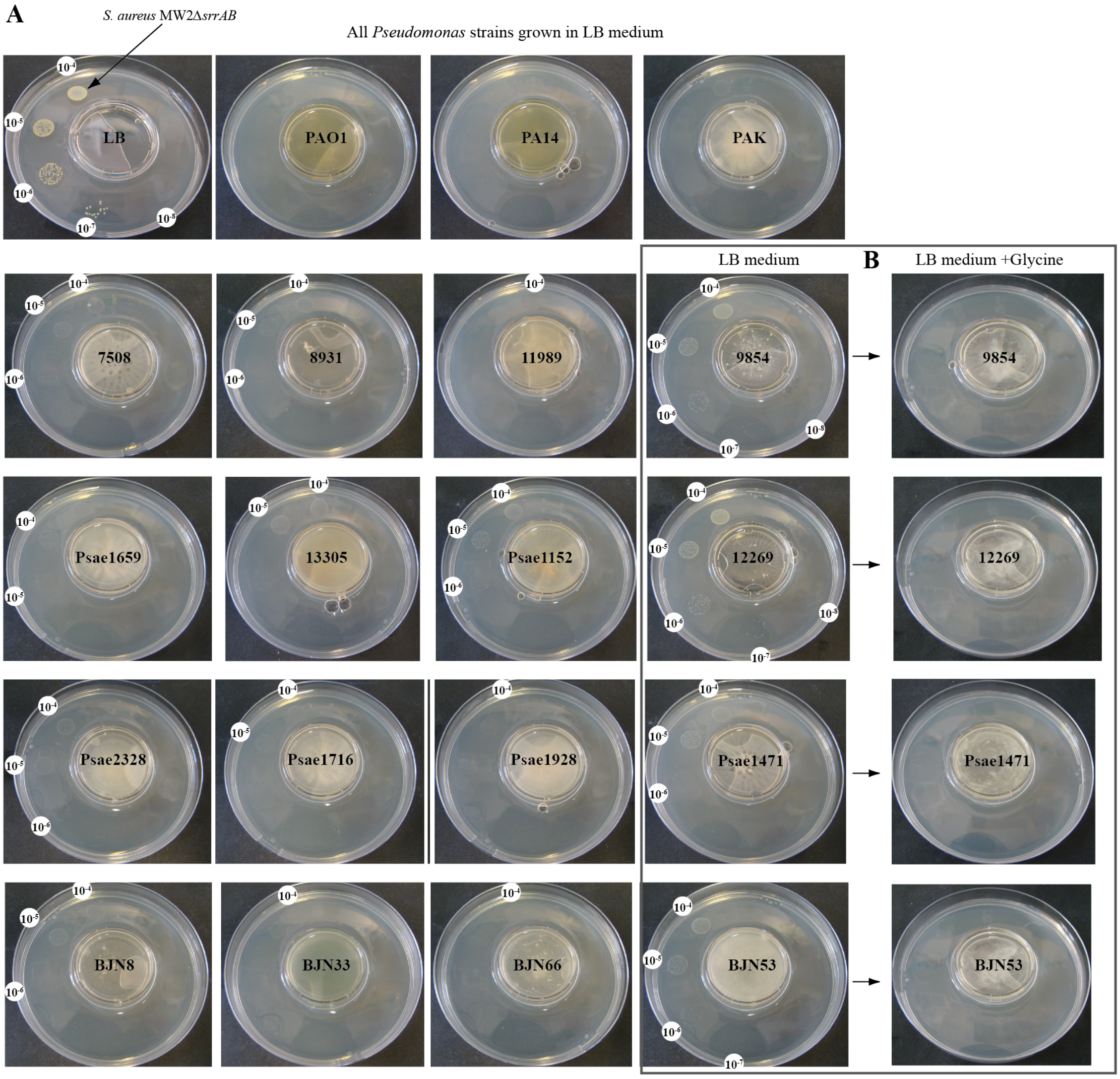


Supplementary figure S3. **Production of biogenic HCN is widespread among *Pseudomonas aeruginosa* strains and clinical isolates. A:** Growth of serial dilutions of *S. aureus* MW2 *srrAB* mutant upon exposure to a panel of laboratory and clinical *P. aeruginosa* strains grown in LB after 24h incubation at 37°C in aerobic conditions, using the 2-petri-dish assay described in Fig S1. **B:** For strains that only partially showed reduced *S. aureus* growth, 0.4% (w/v) glycine was also added in LB medium. Each experiment was performed at least three times. The dilutions were indicated on visible growth spots.

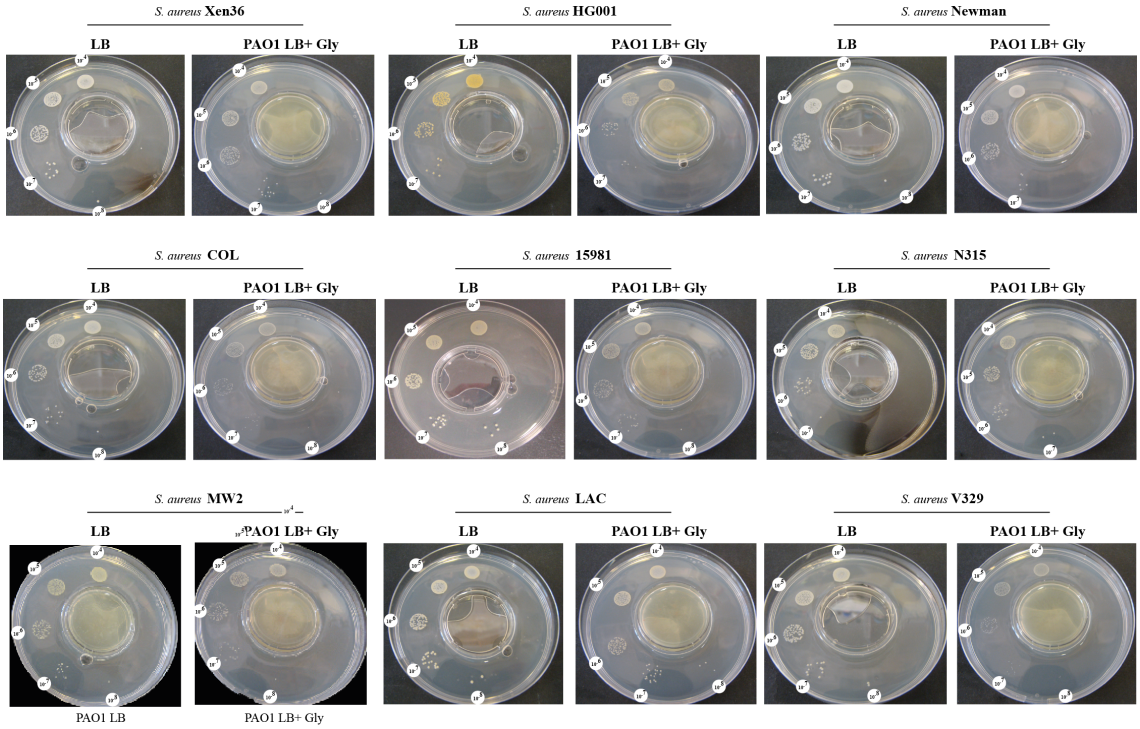


Supplementary figure S4 **Growth inhibition of a panel of *S. aureus* strains upon aerial exposure to *P. aeruginosa* PAO1 culture.** The growth of serial dilution of a panel of *S. aureus* strains upon exposure to *P. aeruginosa* PAO1 cultures in LB supplemented with 0.4% (w/v) glycine after 24h incubation at 37°C in aerobic conditions, using the 2-petri-dish assay as described in Fig S1. Each experiment was performed at least three times.
